# Discrete populations of isotype-switched memory B lymphocytes are maintained in murine spleen and bone marrow

**DOI:** 10.1101/825224

**Authors:** René Riedel, Richard Addo, Marta Ferreira-Gomes, Gitta Anne Heinz, Frederik Heinrich, Jannis Kummer, Victor Greiff, Daniel Schulz, Cora Klaeden, Rebecca Cornelis, Ulrike Menzel, Stefan Kröger, Ulrik Stervbo, Ralf Köhler, Claudia Haftmann, Silvia Kühnel, Katrin Lehmann, Patrick Maschmeyer, Mairi McGrath, Sandra Naundorf, Stefanie Hahne, Özen Sercan-Alp, Francesco Siracusa, Jonathan Stefanowski, Melanie Weber, Kerstin Westendorf, Jakob Zimmermann, Anja E. Hauser, Sai T. Reddy, Pawel Durek, Hyun-Dong Chang, Mir-Farzin Mashreghi, Andreas Radbruch

**Author notes:** Correspondence Bioinfomatics: Pawel Durek, Rheuma-Forschungszentrum Berlin, Charitéplatz 1, 10117 Berlin, Germany. Corresponding Author: Andreas Radbruch or Mir-Farzin Mashreghi, Deutsches Rheuma-Forschungszentrum Berlin, Charitéplatz 1, 10117 Berlin, Germany; or. equal contribution (Richard Addo, Marta Ferreira-Gomes, Gitta Anne Heinz). These authors share senior authorship (Hyun-Dong Chang, Mir-Farzin Mashreghi, Andreas Radbruch).

## Abstract

Here we describe tissue-resident memory B lymphocytes of spleen and bone marrow. Single cell transcriptomes and B cell receptor repertoires identify several exclusive populations of isotype-switched memory B cells (Bsm) in murine spleen and bone marrow, and one interconnected population of 10-20%. A population of marginal zone-like Bsm is located exclusively in the spleen while a novel population of quiescent Bsm is located exclusively in the bone marrow. Cells of two further populations, present in both, spleen and bone marrow, differ in repertoire between the two organs, i.e. are resident as well. Finally, another interconnected population of Bsm of the B1 lineage is present in spleen and bone marrow. In the bone marrow, all Bsm individually dock onto VCAM1+ stromal cells, resting in terms of activation, proliferation and mobility. The discrete B cell memory of bone marrow may be key to rapid secondary humoral responses to systemic antigens.

## Introduction

At present, it is an open question whether, and if so which populations of memory B cells are maintained as circulating cells and/or as resident cells of particular tissues, as has been described for tissue-resident memory T cells^1^ and resident memory plasma cells^2^. Memory B cells expressing antibodies of switched isotype (Bsm) have been identified in different tissues and in the blood, and have been postulated to be part of one uniform, circulating population^3^. In humans, the spleen has been identified as a major hub for circulating *Vaccinia*-specific memory B cells^4^ and splenectomy leads to gradual reduction in numbers of Bsm in the blood to about 50%^5^, suggesting that tissues other than the spleen can also serve as hubs for circulating memory B cells.

Here, we describe a major population of isotype-switched memory B cells in the bone marrow (BM) of inbred and feral mice. Bsm of the BM and spleen of individual mice have significantly different repertoires, demonstrating that overall, they constitute separate compartments. In the BM, Bsm rest in terms of proliferation and individually dock onto VCAM1+/fibronectin+ stromal cells, as described for memory T^6^ and memory plasma cells^7, 8^. Based on their transcriptomes, Bsm of BM could be separated into five different clusters, four of which are also found in the spleen. One cluster is exclusive to BM and represents IgG1+ Bsm expressing little *Cr2* and *Fcer2a*, encoding CD21/35 and CD23, respectively. These Bsm are quiescent in terms of proliferation, transcription and activation, and their B cell receptors show an accumulation of somatic hyper-mutations. Very few if any of these Bsm are found in the spleen. Instead, in spleen an exclusive cluster of Bsm expressing IgG1 or IgG2, *Cr2* and *Fcer2a* is present, with transcriptomes resembling those of marginal zone B cells^9^. Of the four Bsm clusters found in both spleen and BM, two have organ-exclusive repertoires and two have significantly overlapping repertoires. Mutational trajectories link one of those clusters to the clusters exclusive to BM and spleen, respectively. Bsm memory is thus maintained in shared and exclusive compartments in a secondary lymphoid organ, i.e. the spleen, and in the BM, which harbors an exclusive population of quiescent, affinity-matured Bsm.

## Results

### Switched memory B cells are abundant in spleen and bone marrow of mice

Enumeration of CD19+CD38+CD138-GL7-memory B cells expressing IgA, IgG1 or IgG2b, i.e. switched memory B cells (Bsm), in spleen, lymph nodes, BM, Peyers patches and blood of individual mice, revealed that despite a large variability in total cell numbers, most Bsm were located in spleen, BM and lymph nodes (Table 1, Figure S1). In immunized C57BL/6 mice, kept under specific pathogen-free conditions, and in mice obtained from local pet shops, the spleen contained two to three times more Bsm than the BM. In these immunized C57BL/6 mice and pet shop mice, 18-41% of switched Bsm were located in the BM, 9-14% in peripheral lymph nodes and 32-60% in the spleen (Figure S1c, d). Remarkably, the spleens of feral mice (wild mice) were considerably smaller than those of C57BL/6 mice and pet shop mice (Figure S1e) as has been previously reported for feral *M. musculus* d*omesticus* mice^10^. These mice had about equal numbers of Bsm in BM and spleen. While these numbers reflect steady-state conditions in pet shop mice and feral mice, following s.c. injection of NP-KLH with LPS as an adjuvant, in C57BL/6 mice the vast majority of NP-specific Bsm were located in the BM more than 420 days after immunization (Figure 1a), demonstrating that in this particular immune reaction, Bsm can be preferentially maintained in the BM.

**Figure 1:**
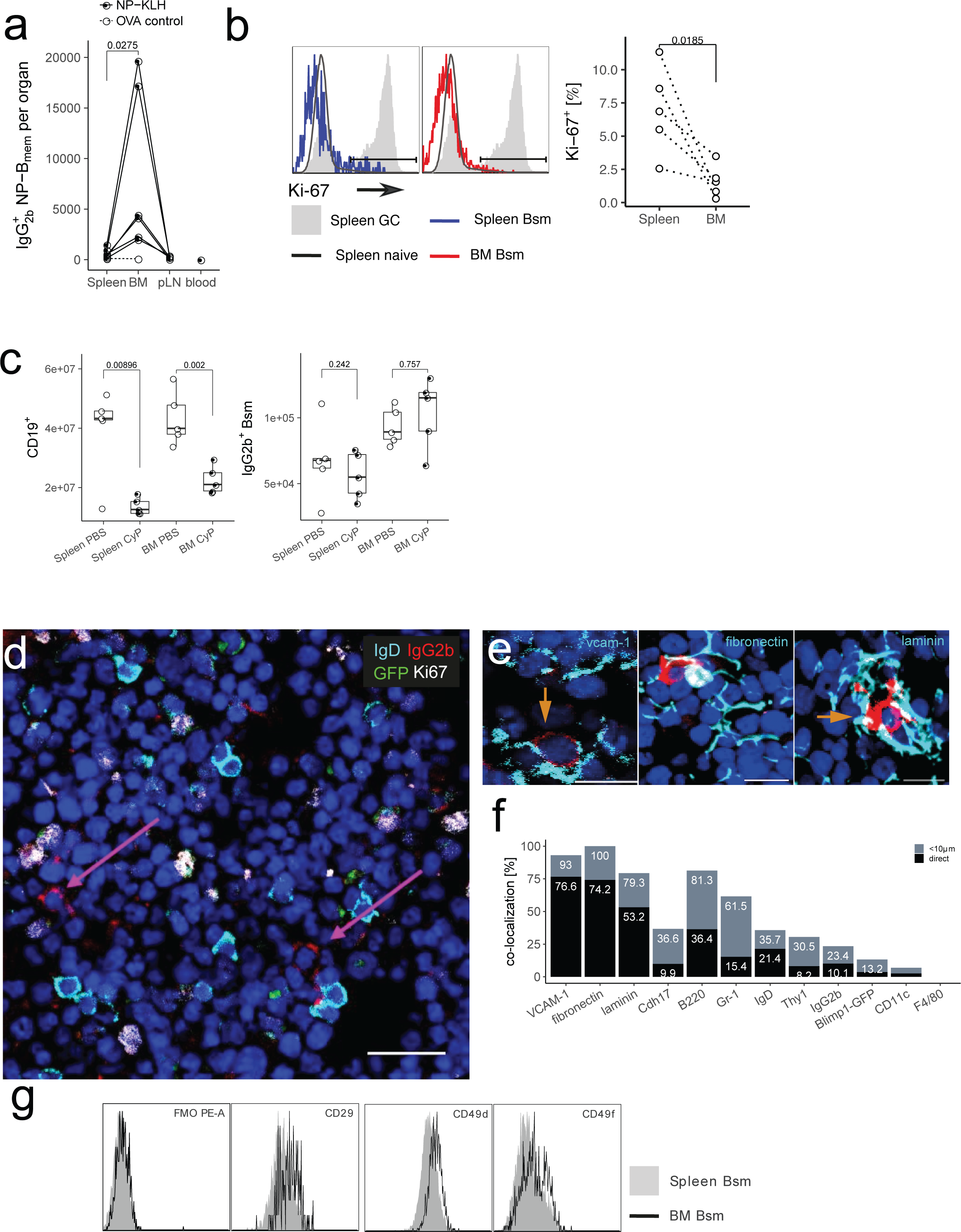
The bone marrow contains a major population of isotype-switched non-proliferating memory B cells. a) Quantification of NP-specific IgG2a/b+ spleen, peripheral lymph nodes, blood and BM memory B cells. C57BL/6 mice were immunized with NP-KLH/LPS SC. Numbers of NP-binding+ IgG2b+ cells in Spleen, BM, blood, and peripheral lymph nodes (pLN) were determined by flow-cytometry on d421 or d426 post immunization; data pooled from 2 independent experiments. OVA ctrl indicates staining controls from mice immunized with the irrelevant antigen ovalbumin (OVA). Gated for IgG2b+CD19+CD38+CD138-GL7-CD11c-IgM-IgD-PI-small lymphocytes. Lines connect samples from one individual, p value of paired t test for spleen and BM samples is indicated. b) Flow-cytometric quantification of Ki-67 expression in IgG2b+ Bsm (IgG2b+CD19+CD38+CD138-GL7-CD11c-IgM-IgD-PI-small lymphocytes) splenic naïve (IgM+IgD+IgG2b-CD19+CD38+CD138-GL7-CD11c-PI-small lymphocytes) and germinal center (GC) (CD19+CD38loGL7+CD11c--PI-lymphocytes) B cells. Indicated are the frequencies of Ki-67+ cells within the subset indicated, data in right graph from 2 independent experiments using pooled cells from 4-20 C57BL/6 mice, p value (paired t test) as indicated. c) Flow-cytometric quantification of CD19+ B cells and IgG2b+ memory B cells in mice treated with Cyclophosphamide (CyP) or untreated controls (PBS) after immunization with 3x NP-CGG/IFA. Anaylsis was performed after 7 days of CyP. IgG2b+ B cells were quantified as IgG2b+CD19+CD38+CD138-GL7-CD11c-IgM-IgD-PI-small lymphocytes, CD19+ B cells as CD19+CD138-PI-lymphocytes, p value (Welch’s test). Representative data shown for one out of two independent experiments. d) IgG2b+ B memory cells (Ki-67-IgD-Blimp1-GFP-) are dispersed as single cells throughout the bone marrow. Arrows indicate IgG2b+DAPI+ cells. Scale bar: 20µm. e) Co-localization of IgG2b+GFP-IgD-IgG2b+ cells (arrows) with mesenchymal stromal cells. Arrows indicate IgG2b+DAPI+ cells. Representative micrograph. Scale bars: 10µm. f) Co-localization of IgG2b+ cells to mesenchymal stromal cells. Graph shows frequency of IgG2b+ cells in direct contact (black) or within 10mm (grey) of a cell stained for the molecule indicated. g) Flow cytometric quantification of surface expression of the VLA-4 and VLA-6 components CD29, CD49d, CD49f in spleen and BM IgG2b+ Bsm. Gated for IgG2b+CD19+CD38+CD138-GL7-CD11c-IgM-IgD-PI-small lymphocytes, histogram plots are representative of three biological replicates.

**Table 1:** Tissue distribution ofisotype-switched B _mem_. Mean values (± SEM) of IgG _1_ ^+^, IgG _2b_ ^+^ and IgA ^+^ CD19 ^+^ CD138 ^−^ CD38 ^+^ CD11c ^−^ GL7 ^−^ IgM^−^ IgD^−^ small lymphocytes; total cell numbers as calculated per organ.

|  | Isotype | Spleen | Bone Marrow | peripheral Lymph Nodes | mesenteric Lymph Nodes | Peyer's Patches | Blood |
| --- | --- | --- | --- | --- | --- | --- | --- |
| C57BL/6 SPF | IgA | 53,033<br>± 11,414 | 35,634<br>± 8,754 | 9,090 ± 3,290 | 4,304 ± 1,398 | 691 ± 560 | 1,976<br>± 455 |
| | IgG1 | 63,317<br>± 21,266 | 24,390<br>± 9,098 | 10,774 ± 3,760 | 9,995 ± 2,127 | 3,737 ± 1,363 | 392<br>± 166 |
| | IgG2b | 283,616<br>± 43,693 | 106,280<br>± 21,839 | 44,472 ± 9,207 | 18,960 ± 9,382 | 9,287 ± 6,027 | 2,113<br>± 580 |
| pet shop | IgA | 61,850<br>± 18,233 | 77,630<br>± 39,883 | 26,300 ± 17,838 | 8,907 ± 1,996 | 13,013 ± 6,946 | 2,130<br>± 590 |
| | IgG1 | 222,179<br>± 59,635 | 148,577<br>± 81,700 | 58,646 ± 17,547 | 36,323 ± 10,256 | 17,609 ± 6,470 | 2,394<br>± 806 |
| | IgG2b | 1,385,442<br>± 390,533 | 425,361<br>± 223,940 | 221,403 ± 42,907 | 147,720 ± 25,912 | 136,631<br>± 44,282 | 9,827<br>± 2,565 |
| wild | IgA | 12,873<br>± 5,428 | 12,904<br>± 3,853 | 3,602 ± 733 | 340 ± 202 | 3,758 ± 1,624 | 2,169<br>± 952 |
| | IgG1 | 55,717<br>± 19,544 | 38,719<br>± 12,991 | 53,537 ± 19,914 | 797 ± 348 | 2,305 ± 784 | 2,877<br>± 713 |
| | IgG2b | 104,652<br>± 37,075 | 73,974<br>± 27,089 | 75,657 ± 28,597 | 1,321 ± 467 | 4,503 ± 1,070 | 7,373<br>± 1,882 |

### Switched memory B cells of the bone marrow are resting in G_0_ of cell cycle

Most Bsm of BM and spleen are resting in terms of proliferation, according to staining with Ki-67 (Figure 1b). Ki-67 was expressed by no more than 9.3% (median: 8.0%) of Bsm in spleen, and 4% (median: 2.6%) of Bsm of BM, i.e. more than 90% of cells in the spleen and more than 95% in the BM were in the G_0_ phase of cell cycle. Accordingly, Bsm of both spleen and BM were refractory to treatment with cyclophosphamide (Fig. 1c), a drug which eliminates proliferating cells. In contrast, the numbers of total CD19^+^ B cells of the spleen and BM were significantly reduced after one week of cyclophospamide treatment.

### In the bone marrow Bsm individually co-localize with VCAM-1^+^ stromal cells

In histological sections of the BM of Blimp1-GFP C57BL/6mice, we identified IgG2b+ Bsm as IgG2b+IgD-Ki-67-nucleated cells. IgG2b+ plasma cells were excluded from analysis according to expression of green fluorescent protein (GFP) under control of the *Prdm1* (Blimp1) promoter (Figures 1d, Figure S1f). IgG2b+ Bsm were dispersed as single cells throughout the BM (Figure 1d). In histological sections 75% of IgG2b+ Bsm were observed in direct contact with cells expressing VCAM-1 and fibronectin (Figure 1e, f), and a further 15-20% of Bsm within 10µm vicinity of such stromal cells (Figure 1f). 53% of the Bsm were directly contacting laminin-expressing stromal cells, and another 26% were in the 10µm vicinity of such cells (Figure 1f).

Contact of IgG2b+ Bsm to VCAM-1+ stromal cells is deterministic, since it is significantly different from random association between the two cell types, as determined by simulation of random co-localization (Figure S1g)^7^. The co-localization of Bsm and stromal cells is in line with expression of VLA4 (CD49d/CD29), a receptor for fibronectin and VCAM-1, and VLA6 (CD49f/CD29), a receptor for laminin^11^, by Bsm (Figure 1g, CD19 staining and cell size shown in Figure S1h). About 10% of Bsm were in direct contact and 26% within 10µm vicinity of cadherin 17 (Cdh17)-expressing stromal cells (Figure 1f). Taken together, Bsm are abundant in BM and spleen, where they rest in terms of proliferation. In the BM, Bsm are individually docked onto stromal cells.

### Switched memory B cells of bone marrow and spleen have distinct Ig repertoires

Comparing the BCR repertoires of Bsm of spleen and BM of individual mice on the level of complementarity-determining region 3 (CDR3) of their immunoglobulin heavy chains, revealed only marginal overlap of CDR3 repertoires between Bsm expressing the same isotype residing in the spleen or BM of individual mice. This is shown in Figure 2 and S2 for IgG1/2+ and IgA+ Bsm of three individual C57BL/6J mice, which were immunized three times with NP-CGG. Biological and technical replicates served to determine how representative the samples were, and to determine reproducibility (Figure S2a). Cosine similarity, a measure to determine the similarity of two groups irrespective of size, was significantly higher for biological replicates (0.65-0.97) than between samples from spleen and BM of each mouse (cosine similarity ∼0.4) (Figure S2b). Moreover, we simulated random overlap between two biological samples by randomly reshuffling the sequences observed, resulting in significantly (P<0.001) higher overlaps (Figure 2b, d) than those observed (Figure 2a, c). Remarkably, several of the most frequently expressed Ig heavy chain V-genes were found predominantly either in the spleen or in the BM, e.g. IGHV1-5 of mice 1 and 3 (Fig. 2e). The segregation of repertoires of Bsm of spleen and BM strongly argues for a tissue-specific compartmentalization of Bsm.

**Figure 2:**
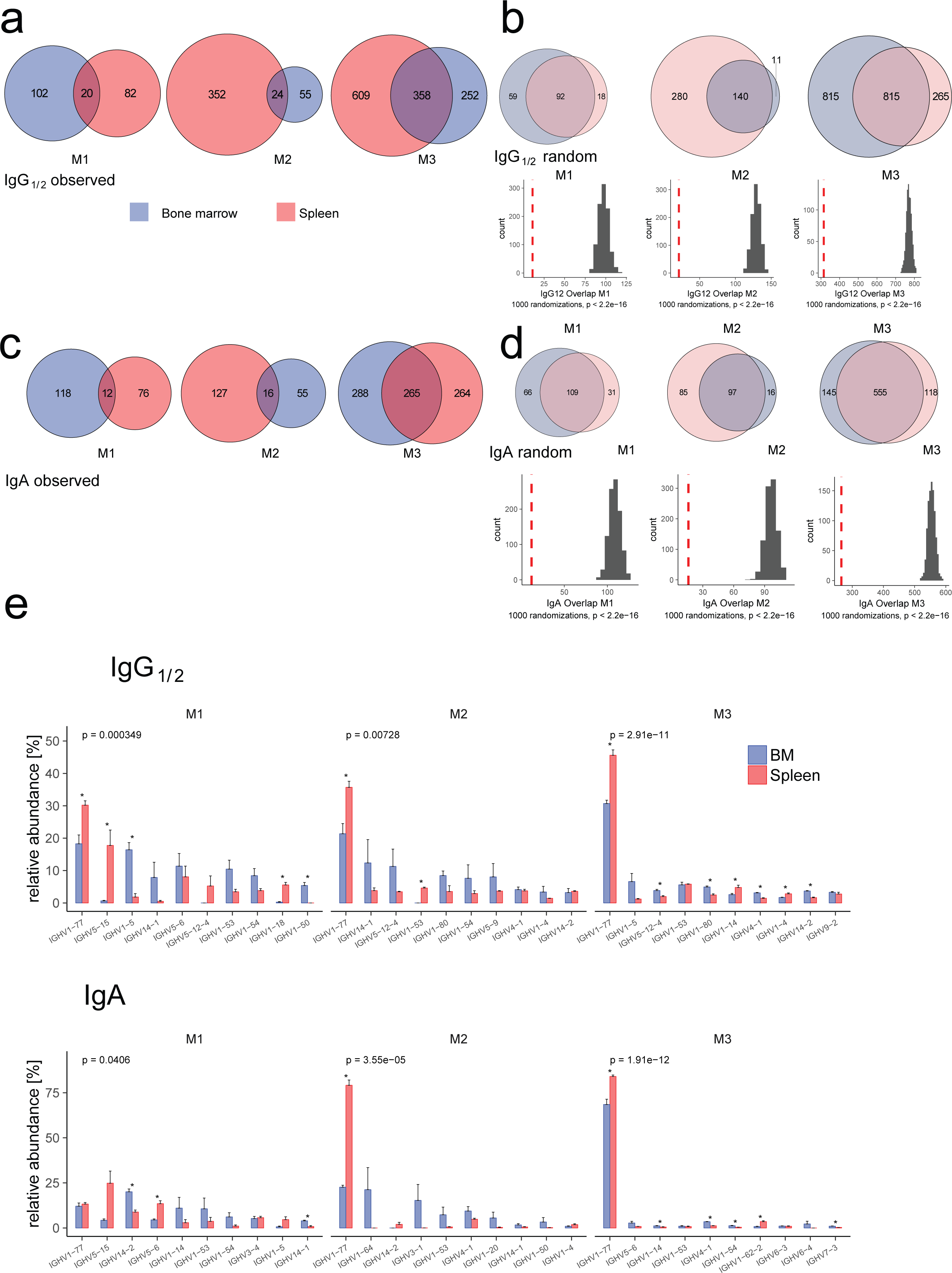
Spleen and bone marrow isotype-switched memory B cells are distinct in Ig heavy chain repertoire. a) and c) Observed overlap between the IgG1/2+ or IgA+ heavy chain CDR3 repertoire of switched memory B cells from spleen and bone marrow or b and d) random distribution (upper panels). Venn diagrams represent clonotype presence in a given sample: numbers indicate clonotypes present in one organ exclusively or in both (overlap). Random distributions (b and d) show median values for 1000 random distributions drawn while retaining the number of clones initially observed in the sample. Histograms (b and d, lower panels) represent number of clonotypes in spleen and BM in 1000 randomized distributions of observed clonotypes to spleen and BM, dashed red line indicates experimentally observed number of clonotypes present in both spleen and BM. P value of one-sided t test for difference of randomized overlap against observed. e) VH gene recruitment to spleen and bone marrow IgG1/2+ (upper panel) and IgA+ (lower panel) memory B cells, represented as frequency of a particular VH gene among total CDR3s per organ. Bars show relative abundance of the 10 most abundant VH genes, error bars indicate SEM. Significance of difference in VH gene distribution to Spleen and BM assessed by MANOVA, p values corrected for multiple testing (Benjamini-Hochberg), * indicates significant difference in means for a particular VH gene (Welch’s test). BM: bone marrow, M1-M3: replicate samples of three C57BL/6 mice immunized 3x NP-CGG/IFA. Only clones consistently found in technical replicates were considered.

### Transcriptional heterogeneity of Bsm in bone marrow and spleen

In order to resolve the heterogeneity of Bsm within as well as between spleen and BM, we analyzed single-cell transcriptomes of Bsm isolated from both organs of two individual mice. Bsm from C57BL/6J mice immunized three times with NP-CGG/incomplete Freund’s adjuvant (IFA) i.p. 60 days prior to analysis were enriched by MACS with CD19 microbeads (Miltenyi Biotech) and isolated by FACS as IgG1+IgG2b+CD19+CD38+GL7-CD138-IgM-IgD-cells. In total, 6047 Bsm from the spleen and 4164 from the BM were subsequently sequenced using 10X Genomics-based droplet sequencing. An average of 42,407 reads per cell defined a median number of 1629 transcribed genes per cell, corresponding to a sequencing saturation of approximately 70%. For 4,754 Bsm from spleen and 2,947 from BM, we also determined the sequences of their antibody heavy and light chains. In line with the isolation protocol and degree of transcriptome saturation, most cells expressed the B cell marker genes *Cd19* (86% of all cells), *Cd38* (36% of all cells)*, Pax5* (38% of all cells) and *Ptprc* (CD45, 80% of all cells) (Figure S3a). Transcription of *Sdc1* (CD138, <0.1%, a surface protein expressed on plasma blasts and plasma cells) and *Bcl6* (2.5%, a transcription factor characteristic for germinal center B cells) was not detectable overall (Figure S3a). According to their transcriptomes, Bsm robustly clustered into 6 different populations (Figure 3a). Bsm of clusters I, II, III and VI were present in both the spleen and BM of the two mice at varying frequencies. Bsm of cluster IV were almost exclusively located in the spleen, while Bsm of cluster V were virtually exclusively located in the BM (Figure 3b). While clusters III (91% IgG1, 9% IgG2b/c) and V (94% IgG1, 6% IgG2b/c) were predominantly cells of IgG1 isotype, cluster VI was highly enriched for cells of IgG2b/2c isotype (4% IgG1, 96% IgG2b/c). Cluster I (84% IgG1, 16% IgG2b/c), II (36% IgG1, 64% IgG2b/c) and IV (71% IgG1, 29% IgG2b/c) consisted of cells of mixed isotypes (Figure 3c).

**Figure 3:**
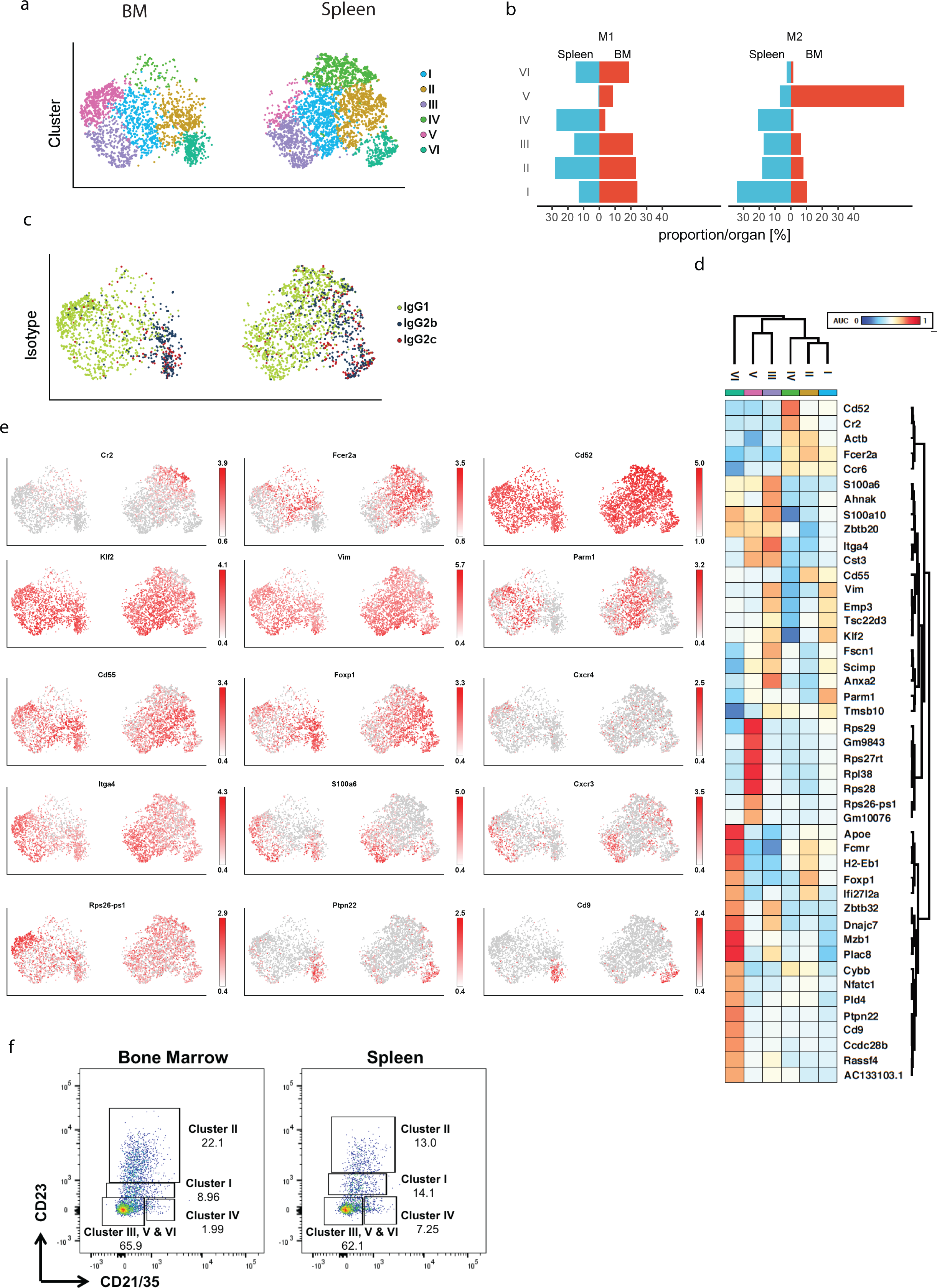
Heterogeneity of switched memory B cells is differentially represented in spleen and bone marrow. Cells for single cell sequencing were FACSorted as IgG-expressing CD19+CD38+CD138-GL7-small lymphocytes. a) Six transcriptionally defined clusters were identified by shared nearest neighbor (SNN) modularity optimization based clustering algorithm mapped to tSNE representation of spleen and BM cells. tSNE coordinates and clustering was computed for 4754 from spleen and 2947 from BM cells, presentation is separated by organ. b) Percentage of cells per cluster in each organ by mouse. c) Distribution of IgG subclass mapped on tSNE. d) Signature genes for each cluster, area under curve (AUC) of responder-operator characteristics (ROC) of >0.7. e) Distribution of transcription levels for representative genes mapped on tSNE. Cells for single cell sequencing were FACSorted as IgG-expressing CD19+CD38+CD138-GL7-small lymphocytes. f) Flow-cytometric evaluation of expression of representative markers to identify clusters observed by transcriptional profiles in IgG+ memory B cells (IgG1+ or IgG2b+CD19+CD38+IgM-IgD-CD138-GL7-CD93-Zombie Aqua-small lymphocytes). Dotplots represent one of three mice.

Signature genes characteristic for the different clusters, with area under curve (AUC) of responder-operator characteristics (ROC) of >0.7, are displayed in Figure 3d. Bsm of Cluster I were characterized by low levels of *Cr2* (CD21) (14% of cells in cluster 1) and intermediate levels of *Fcer2a* (CD23) (52% of cluster I cells) (Figure 3d, 3e). 95% expressed the transcription factor *Krueppel-like factor 2* (*Klf2*), 93% the cytoskeletal protein *Vimentin* (*Vim*)^12^ and 62% the golgi/endosomal-associated gene *Prostate androgen-regulated mucin-like protein 1 (Parm1*)^13^. In summary, Bsm of cluster I resemble transitional memory B cells^14, 15^. 63% of Bsm of cluster II expressed high level of *Fcer2a* and intermediate levels of *Cr2* (24% of cluster II Bsm). Cells of cluster II also expressed the complement decay-accelerating factor *Cd55*^16^ in 75% of the cases and 77% expressed the transcription factor *Foxp1* (*Forkhead Box P1*)^17^. This gene expression pattern suggests that Bsm of cluster II are follicular memory B cells^14^. Bsm of cluster III resemble “age-associated memory B cells” (ABC), in that they expressed low levels of *Cr2* (3%) and *Fcer2a* (15%). We detected high levels of the integrin gene *Itga4* (CD49d) in 91%, *Itgam* (CD11b) in 4% (Figure S3b) and the chemokine receptor gene *CXCR3* in 47% of cluster III Bsm^18^. In addition, 62% of cluster III Bsm expressed the gene encoding the calcium binding protein *S100a6*^19^. However, they lacked the expression of *Itgax*, the gene encoding for CD11c (Figure S3b), which has been associated with ABC (reviewed in Rubtsova et al.^18^). Bsm of cluster IV, residing exclusively in the spleen, resemble marginal zone B cells in that 53% of them expressed high levels of *Cr2* and all of them the gene *Cd52*, and 59% expressed intermediate levels of *Fcer2a*^9, 14^. Cluster V, found exclusively in the BM, is comprised of Bsm expressing low levels of *Cr2,* and *Fcer2a* in 6% and 24% of Cluster V Bsm, respectively. In contrast, they expressed relatively high levels of various genes encoding for ribosomal proteins, such as *Rps26-ps1* (76% of cluster V Bsm). 10% of cluster V Bsm also expressed the gene *Cxcr4,* encoding a chemokine receptor associated with BM homing^20^. Bsm of cluster VI resemble B1 cells, in that they are characterized by expression of the scavenger receptor *Cd5*^21^in 8% (Figure S3b), the Protein tyrosine phosphatase 22 gene *Ptpn22* associated with hypo-responsiveness of B cells in the context of autoimmunity^22^ in 56% and the gene encoding the tetraspanin *Cd9* expressed on regulatory B cells^23^ in 49% of the cells. The expression of *Cr2* and *Fcer2a* as determined by single cell transcriptomics was verified cytometrically on the protein level (Figure 3f). Staining of CD21/35 and CD23 was sufficient to identify Bsm of clusters I, II and IV in spleen and BM. Discrimination between Bsm of clusters III, V and VI would require additional markers.

In addition to the specific gene expression signatures (Fig. 3), Bsm of the clusters differed in the expression of gene sets indicating physiological activities. Single cell gene set enrichment analysis of expression of the Reactome^24^ and Gene ontology (GO)^25–27^ gene sets revealed that Bsm of both organs express gene sets associated with oxidative phosphorylation (Reactome: TCA/electron transport) and integrin signaling (Reactome: Integrin cell surface interactions) (Figure S3c), but not gene sets associated with glycolysis or RNA polymerase II-mediated transcription. Confirming their quiescence, Bsm of cluster V, located exclusively in the BM, did not express gene sets associated with BCR receptor signaling, DNA synthesis or regulation of cell cycle progression. Conversely, many Bsm of cluster IV, located only in the spleen, expressed these gene sets. In addition, these Bsm were exclusively expressing a gene set indicating cognate activation. Bsm of cluster III were exclusively enriched for a gene set associated with lymphocyte migration (GO term lymphocyte migration).

### BCR repertoires and trajectories of spleen and bone marrow Bsm clusters

The comparison of the BCR repertoires (paired Ig heavy and light chain) of the Bsm clusters I, II, and IV showed exclusive repertoires, as determined by simulation of randomized distribution of BCR sequences to the different clusters (Figure 4a), confirming the results obtained for other mice and the bulk analysis of their Bsm of spleen and BM (Figures 2 and S2). A significant overlap of repertoires was found between Bsm of cluster VI from the spleen and BM. The same was true for Bsm of clusters III and V, which showed a significant overlap of their BCR repertoires between clusters as well as between those of spleen and BM. Together these results imply that those Bsm clusters were either interconnected in their generation and/or maintenance. Bsm with a high-degree of somatic hyper-mutation (>1% mutation in framework (FR)1-3) were more frequent in clusters I, III, IV and V (Figure 4b) than in clusters II and VI. Interestingly, the genes encoding for CD80 (*Cd80*) and CD273 (*Pdcd1lg2*), which have been described to correlate with an increase in somatic hypermutations^28^ were significantly enriched in those clusters (Chi-square with Yates’ correction comparing distribution of either CD80 or CD273 expressing cells in clusters I,III,IV and V as well as in cluster II and VI, significance level of p<0.01). In contrast, the number of *Nt5e* (CD73) expressing cells was reduced in clusters I, III, IV and V, as compared to random distribution of transcription (Figure 4b). Trajectories based on somatic hyper-mutation of Bsm BCR heavy and light chain identified cluster III as the cluster from which clones with higher mutation counts would have originated. A total of 187 significant clones (significantly overrepresented compared to randomization) were observed in other clusters originating from either spleen or BM cluster III. The frequencies of transitions, i.e. presence in more than one cluster, are indicated in Figure 4c, taking into account the population size of each cluster in each organ. A trajectory was considered significant if a p-value <0.01was observed (indicated in red, Figure 4a, c), compared to random distribution of mutated BCR clones into different clusters. Figure 4d indicates the direction of mutational trajectories, and the number of clones involved. Of the 187 clones of cluster III, 55 clones were represented with additional mutations to the very same cluster of the other organ. 45 of the cluster III clones from spleen or BM were represented with additional mutations in cluster V of the BM, 23 clones were among the rare cells of cluster V of the spleen. 40 clones of either spleen or BM cluster III were also identified in cluster I of the BM, with additional mutations. There was a significant exchange between splenic and BM cluster V (14 clones) as well as a minimal but significant exchange between the splenic and BM cluster VI with either splenic or BM cluster II (8 clones). Additionally, we were able to detect a significant exchange between BM cluster III and the spleen-exclusive cluster IV (24 clones), as well as between splenic and BM cluster III and splenic cluster V (23 clones).

**Figure 4:**
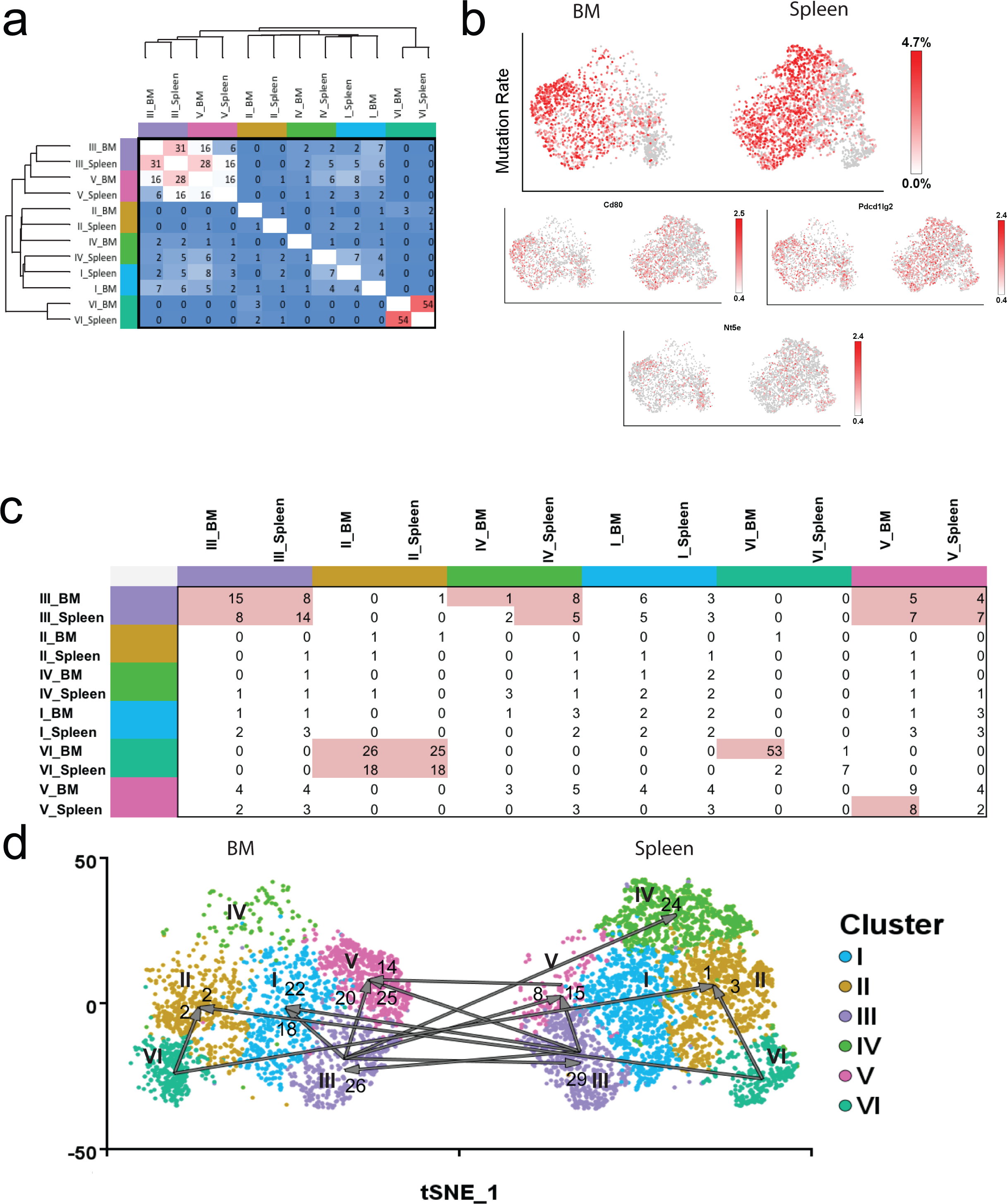
a) Percentage of shared clones between clusters of spleen and BM. Clones were defined according to IgG heavy and light chain sequences. Overlap significantly higher than expected for random overlap (p<0.05) is labelled by red shading, white represents expected random overlap, and blue shading an overlap lower as random overlap, i.e exclusive repertoires. b) Mutation rates (percent of nucleotide sequence; upper panel) and CD80 (*Cd80*), CD73 (*Nt5e*) and PD-Ligand 2 (*Pdcd1lg2*) gene expression of individual cells. Cells for single cell sequencing were isolated by FACS as IgG+CD19+CD38+CD138-GL7-small lymphocytes.c) Frequencies of Bsm clones represented with higher mutation rates in clusters listed in rows, as compared to clusters listed in columns. Significant presence of clone members with additional mutations as compared to random distribution (p<0.01) is shaded in red. d) Trajectories are based on accumulation of mutations within clones. Represented are significant trajectories (p<0.01). Arabic numbers indicate numbers of clones with additional mutations.

In summary, mutational trajectories link cluster II to cluster VI and cluster III to cluster V, and to lesser degree to cluster I. Together these data suggest that Bsm of cluster III of spleen and BM represent one interconnected compartment.

## Discussion

In secondary immune responses, memory B lymphocytes provide rapid immune protection, with antibodies of enhanced affinity and adapted function, which is reflected by their switched isotype. While the generation of memory B lymphocytes and their role in secondary immune responses has been subject of intense research, little is known about their maintenance over time in the apparent absence of antigen. For memory T lymphocytes, relatively new research has unveiled a highly structured organization of their maintenance, describing populations of circulating and tissue-resident memory T cells (Trm)^2^. Trm have been described for nearly all organs investigated, including the BM^6, 7, 29^. In contrast, the organization of B cell memory has been less clear. Memory B lymphocytes expressing antibodies of switched isotype (Bsm) can be detected in many organs^3^. Whether they constitute one interconnected population of circulating cells, or whether they are compartmentalized into resident populations, and according to which mechanisms, have remained enigmatic.

By comparing the antibody repertoires of Bsm of spleen and BM of individual mice, we show that both organs host prominent populations of exclusive, i.e. resident and tissue-specific, as well as smaller populations of connected Bsm, with overlapping repertoires. Transcriptomes of individual Bsm from murine spleen and BM define 6 clusters, 4 of which are present in both, spleen and BM. One is only found in spleen and another one only in the BM.

Cluster I, a major population of Bsm in both, spleen and BM, consists of Bsm expressing low levels of CD21 and intermediate levels of CD23. This combination is characteristic for transitional, immature B cells (T1/2), which are believed to represent a developmental stage of negative selection against autoreactivity in BM (T1) and spleen (T2)^14^. Classical transitional B lymphocytes express IgM and IgD, and so far^30^, isotype-switched transitional B cells have not been described. However, the antigen-receptor repertoires of Bsm of cluster I of spleen and Bm differ, as well as being distinct from those of all other Bsm, indicating that Bsm of cluster I are resident cells, selected in exclusive immune reactions. It should be noted that those repertoires show a high degree of somatic hyper-mutation, indicating an extensive selection process.

Bsm of cluster II express intermediate levels of CD21 and high levels of CD23, a pattern reminiscent of follicular B cells, generated in germinal center reactions^14^. Their antigen receptor repertoires are different from cells from spleen and BM, and also different from all other Bsm. Again indicating that Bsm of BM and spleen are resident cells, generated in different immune reactions and most likely in germinal centers^31^. While their antibodies show somatic hyper-mutation, its extent is lower than that of most other Bsm, except those of cluster VI.

Cluster III consists of Bsm expressing low levels of CD21 and CD23, and accumulated cells expressing CD11b and CXCR3, all markers of “aged” memory B cells^18^. T-bet, another marker gene of “aged” memory B cells, was not quantitatively detectable in the present analysis either due to the sequencing depth or low expression of this transcription factor per cell. “Aged” Bsm have been described to be generated in germinal centers by combined activation of B cells through TLR and antigen receptors^32–34^. Bsm of cluster III express highly mutated antibodies, and their repertoires overlap significantly between spleen and BM. Moreover, their repertoires also overlap with Bsm of cluster V in BM, and the few cells of cluster V in the spleen.

Interestingly, mutational trajectories indicate that Bsm of cluster III are the precursors of those of cluster V. Mutational trajectories also link Bsm of cluster III of spleen and BM, in both directions, indicating that those clusters either would have been connected during establishment, and/or are still connected during maintenance of memory, i.e. represent circulating Bsm. In line with this, Bsm of cluster III are exclusively expressing a gene set associated with lymphocyte migration.

Bsm of cluster IV are found almost exclusively in the spleen. They express high levels of CD21 and intermediate levels of CD23. This, together with their transcriptional signature classify them as marginal zone-like memory B cells^9, 14^. Bsm of marginal zones have been described before^35^. Here we show, that their antigen receptors are highly mutated and that their repertoire is different from the repertoire of Bsm of all other clusters, indicating that they are a resident population, generated in exclusive immune reactions.

An exclusive BM population of Bsm is seen in cluster V. Bsm of cluster V express low levels of CD21 and CD23. They are quiescent, as reflected by their lack of expression of gene sets associated with BCR receptor signaling, DNA synthesis or regulation of cell cycle progression. In relation to other genes, they express high levels of housekeeping genes encoding ribosomal proteins^36^. Their antigen receptor repertoire is different from that of Bsm of clusters I, II, IV and VI, but shows considerable overlap with Bsm of cluster III (see above), which according to mutational trajectories, qualify as their precursor cells. Interestingly, different from Bsm of cluster III, Bsm of cluster V, however, do not express genes associated with migration.

Finally, Bsm of cluster VI, present in both spleen and BM, are exclusively IgG2 expressing cells. They express CD5 and CD9, markers of cells of the B1 lineage ^21^. Repertoires of Bsm from spleen and BM overlap significantly, showing that these populations have been connected in development. It should be noted, however, that Bsm of cluster VI do not express genes associated with lymphocyte migration in the maintenance phase of memory.

In summary, the clustering of Bsm according to their transcriptomes reveals an unforeseen heterogeneity, with populations so far not described until now (clusters I, V), or poorly characterized (cluster III, IV). The vast majority of these cells, those of clusters I, II, IV, and V, VI are apparently permanent residents of their tissue, since they have exclusive antigen receptor repertoires and/or exclusive location, and they do not express genes associated with lymphocyte migration. 10 to 20% of Bsm of spleen and BM, those of cluster III, resemble “aged” memory B cells and qualify as circulating memory B cells, with largely overlapping repertoires of spleen and BM, and expression of genes associated with migration. Thus, as described for memory T lymphocytes^2^, a significant proportion of Bsm of BM are resident.

Moreover, Bsm are apparently maintained in the BM like memory T and memory plasma cells^2^, where they individually dock onto mesenchymal stromal cells, expressing VCAM1, which provide a niche for their maintenance. We did not observe a preferential colocalization to Cadherin-17 expressing stromal cells of the BM, a colocalization that has been reported relevant for Bsm of the spleen^37^. Instead, we observed a co-localization of Bsm of the BM to laminin expressing stromal cells. This is in striking homology to IgG-secreting memory plasma cells, which in the BM, but not in the spleen, require laminin-beta1 for their maintenance^38^.

Resident Bsm of secondary lymphoid organs, e.g. the spleen, are able to participate in secondary immune reactions, inside or outside^39, 40^ of germinal centers. However, the raison d’être of resident Bsm of the BM is less clear at first glance. For resident memory T cells of the BM, we have demonstrated their effective reactivation in the BM in “immune clusters” ^41^consisting of reactivated memory T cells and antigen-presenting cells, including B cells^42^.

Whether or not Bsm of the BM are reactivated in a similar way, remains to be shown. Assuming a similar reactivation, this would be an efficient and fast way to secure rapid generation of new (memory) plasma cells and secreted antibodies against blood-borne antigens, to provide enhanced local and systemic protection. The contribution of BM-resident Bsm to secondary germinal center reactions in secondary lymphoid organs remains to be demonstrated, and it would require their mobilization and emigration from the BM. In steady state, only Bsm of cluster III have the transcriptional potential to do so.

## Methods

### Mice

C57BL/6J/N mice were purchased from Charles River (Sulzfeld, Germany) or Janvier Labs (Le Genest-Saint-Isle, France). Mice expressing GFP under the control of the *Prdm1* promoter (Blimp1-GFP)^43^ were bred at the DRFZ animal facility. C57BL/6 and Blimp1-GFP mice were housed under specific pathogen-free conditions. Pet mice were obtained as adult animals at pet shops in Berlin. Feral mice were caught as free-living animals at non-residential farm buildings in Altlandsberg, Brandenburg, Germany. Live traps were set up and controlled by a trained veterinarian. Mice were anesthetized with isofluran prior to sacrifice. All animal experiments were performed according to institutional guidelines and licensed under German animal protection regulations.

### Immunizations and cyclophosphamide administration

Mice aged 8–12 weeks were immunized s.c. with 100μg NP-KLH 10μg LPS (*E. coli*, InvivoGen). For boost immunizations 10μg NP-KLH without adjuvant was used. Alternatively, mice aged 8-12 weeks were immunized three times with 100μg NP-CGG in IFA i.p. at 21 days interval. For further quantification of Bsm in C57BL/6 laboratory animals, in two experiments mice were infected with 2×10^5^ plaque-forming units Armstrong strain of lymphocytic choriomeningitis virus (LCMV) i.p. or, alternatively, infected intravenously with 10^6^ colony-forming units of attenuated *Salmonella enterica* serovar typhimurium strain SL7207.

Three times NP-CGG/ IFA-immunized (at 21 days interval) mice were i.v. administered for one week with either Cyclophoshamide (CyP, 50mg/kg) or PBS starting 30 days after the last immunization. Analysis was performed on day 3 after the last CyP injection.

### Cytometry and cell sorting

In order to stain lymphocytes for multicolor flow cytometry, cells were resuspended in PBS with 0.5% BSA and 5µg per ml Fcγ receptor IIB-blocking antibody (DRFZ, clone 2.4G2) or 10µL of FcR Blocking Reagent (Miltenyi) per 10^7^ cells at up to 5×10^8^ cells per ml. Cell separation by FACS was done at BD FACSAria II. For cytometric analyses Canto II, Fortessa, Symphony (BD) or MACSQuant (Miltenyi) machines were used. Data were analyzed by FlowJo (version 9, TreeStar).

B cells were enriched by magnetic separation using anti-mouse CD19 microbeads (Miltenyi) following incubation of the cells with Fcγ receptor IIB-blocking antibody.

Antibodies against following murine antigens were used: Ki-67 (B56, BD Biosciences), CD11c (N418, DRFZ), CD19 (1D3, DRFZ and 6D5, BioLegend), CD38 (90, BioLegend), CD138 (281-2, BD Biosciences), GL7 (GL7, DRFZ), IgA (C10-3, BD Biosciences), IgD (11.26c, DRFZ), IgG_1_ (A85-1, BD Biosciences and X-56, Miltenyi), IgG2a/b (R2-40, BD Biosciences and X-57, Miltenyi), IgG_2b_ (A95-1, BD Biosciences and MRG2b-85, BioLegend), IgM (M41, DRFZ and RMM-1, BioLegend), CD93 (AA4.1, BioLegend and DRFZ), CD21/35 (7G6, DRFZ and 7E9, BioLegend), CD23 (B3B4, BioLegend), CD29 (HMß1-1, Miltenyi), CD49d (R1-2, BioLegend), CD49f (GoH3, eBioscience), CD62L (MEL-14, DRFZ). Dead cells were excluded using PI staining or the Zombie Aqua Fixable Viability Kit (BioLegend). Flow cytometric measurements and cell sorting were done according to standards defined in the guidelines to flow cytometry and cell sorting in immunological studies^44^.

### Single cell RNA-sequencing and single cell BCR repertoire analysis

Single-cell suspensions from BM of 3x NP-CGG immunized C57BL/6 mice were prepared and CD19+ cells were enriched by magnetic cell sorting using anti-CD19 microbeads (Miltenyi Biotech). *Ex vivo* IgG1+/IgG2b+CD19+CD38+GL7-CD138-IgM-IgD-memory B cells were isolated by FACS (Influx cell sorter (BD Bioscience)) and applied to the 10X Genomics platform using the Single Cell 5’ Library & Gel Bead Kit (10x Genomics) following the manufacturer’s instructions. The amplified cDNA was used for simultaneous 5’ gene expression (GEX) and murine BCR library preparation. BCR transcripts were amplified by Chromium Single Cell V(D)J Enrichment Kit for murine B cells (10x Genomics). Upon adapter ligation and index PCR, the quality of the obtained cDNA library was assessed by Qubit quantification, Bioanalyzer fragment analysis (HS DNA Kit, Agilent) and KAPA library quantification qPCR (Roche). The sequencing was performed on a NextSeq500 device (Illumina) using either a High Output v2 Kit (150 cycles) with the recommended sequencing conditions (read1: 26nt, read2: 98nt, index1: 8 nt, index2: n.a.) or a Mid Output v2 Kit (300 cycles for BCR repertoire analysis (read1: 150nt, read2: 150nt, index1: 8nt, index2: n.a., 20% PhiX spike-in).

### Single cell RNA-sequencing and B cell receptor repertoire analysis

Raw sequencing data for single cell transcriptome analysis were processed using cellranger-2.1.1. *mkfastq* and count commands with default parameter settings, the genome reference: refdata-cellranger-mm10-1.2.0 and expected-cells=3000. Raw sequencing data for single cell immune profiling (B cell receptor sequences (BCR) for IgG heavy and light chain) were processed using cellranger-3.0.2. *mkfastq* and *vdj* commands with default parameter settings and the vdj-reference: *refdata-cellranger-vdj_GRCm38_alts_ensembl-mouse-2.2.0.* Only cells, which appeared in both, the single cell transcriptome as well as immune receptor profiling were further analyzed. The high-confidence contig sequences of the isotype-switched memory B cells were reanalyzed using HighV-QUEST at IMGT web portal for immunoglobulin (IMGT) to retrieve the V-, J- and D-genes as nucleotide and amino acid CDR3 sequence^45^. IMGT-gapped-nt-sequences, V-REGION-mutation-and-AA-change-table as well as nt-mutation-statistics were used to determine the corresponding gapped germline FR1, CDR1, FR2, CDR2, FR3 sequences as well as estimated mutation counts in the FR1-FR3 region. The most abundant contig for the heavy and light BCR chain was assigned to the corresponding cell in the single cell transcriptome analysis. Cells with incomplete heavy and light chain annotation were removed from further analysis. In order to simulate random distribution and determine the significance of observed BCR distribution within the transcriptomic clusters of Bsm in the spleen and BM of the individual Bsm, their BCR were randomly reshuffled 1000 times. For the mutation rates, the sum of the estimated mutation counts in the light and heavy BCR chain in FR1-FR3 regions was normalized by the length of the corresponding FR1-FR3 sequences. The overlap of the BCR-repertoire was computed using the VDJ-gene configuration and the nucleotide sequence of the CDR3-Region for assessing similarity of the BCR and the minimal fraction of common BCR in one of the compared clusters. The analysis was repeated for 1000 random annotations of the BCRs. Overlap was defined as significantly higher or lower than expected if occurred in less than 5 or more than 95 of random randomizations. Clustering was performed using Ward2 and the Manhattan distance. For the BCR trajectory analysis, clonal families were clustered by the identical isotype annotation, VDJ-gene usage, gapped germline FR1-FR3 sequence and nucleotide CDR3 sequence length of the heavy and light chain. For each clonal family a lineage tree was computed using GLaMST with concatenated FR1-FR3 sequences of the heavy and light chain and the germline sequence as root as input^46^. An accumulation of mutations was assumed between a FR1-FR3 sequences and the first “ancestor” sequence found in the sample. For each cluster, the proportion of ancestor sequences (member of a clonal family, which have descendants with higher mutations rates) was computed. A proportion was defined as significant if occurred in less than 50 of 1000 randomizations (p<0.05). For graphical projections of trajectories, clusters were connected if a descendant-ancestor sequence relationship was found between these clusters. Significant connections between two clusters were assumed if they occured in less than 10 randomizations. The single cell transcriptome data was further analyzed using R 3.5.0., and Seurat R package 2.3.4. In particular, gene expression levels were normalized and the detection of variable genes, t-distributed stochastic neighbor embedding (tSNE) as well as clustering by shared nearest neighbor (SNN) modularity optimization was performed by using 5 out of 30 principle components for tSNE and a modularity of 0.6 for clustering. The responder-operator characteristics (ROC) for each variable gene was computed for each cluster against the remaining cells. The ROC-values above 0.7 were clustered using Ward2 linkage, Euclidian distance. Single cell transcriptome data as well as immune profiling data data discussed in this publication are available at gene expression omnibus (GEO) under accession number XXX (Link).

### Ig sequencing and repertoire analysis

Cells from BM (tibiae, femurae and pelvic bones) and spleen of three times NP-CGG immunized-mice were isolated and Bsm were magnetically enriched using the Memory B cell Isolation Kit (130-095-838 Miltenyi). Bsm were further enriched by FACSorting of CD19+CD38+CD138-CD11c-GL7-IgM-IgD-small lymphocytes. Sorted cells of Mouse 1 and 2 were split into two equal aliquots (samples A, B) for BM and spleen samples as cellular (biological) replicates^47^. Sorted cells of Mouse 3 were split into 4 samples (A, B, C, D).

Biological replicates were processed independently from this point on. Cell samples were lysed in RNA lysis buffer (R1060-1-50, Zymo Research) and stored at -80°C. Total RNA was extracted from samples using the ZR RNA Miniprep Kit (Zymo Research) according to the manufacturer’s protocol (Catalog nos. R1064 & R1065). Isolated RNA was split and library preparation was performed in technical duplicates.

First-strand cDNA was synthesized with SMARTScribe Reverse Transcriptase (Clontech) using total RNA, a cDNA synthesis primer mix (mIgG12ab_r1(KKACAGTCACTGAGCTGCT), mIgG3_r (GTACAGTCACCAAGCTGCT), mIgA_r (CCAGGTCACATTCATCGTG) by Metabion international AG) and a 5’ – template-switch adaptor with unique molecular identifiers (UMI) (SmartNNNa (AAGCAGUGGTAUCAACGCAGAGUNNNNUNNNNUNNNNUCTT(rG)4)) according to the protocol “high-quality full length immunoglobulin profiling with unique molecular barcoding” by the Chudakov lab^48^. cDNA was purified with MinElute PCR purification Kit (Qiagen) and eluted in 10 µL 70°C nuclease-free H_2_O (Qiagen).

The first PCR was performed according to the protocol of the Chudakov lab^48^ using a Step-out primer M1SS (AAGCAGTGGTATCAACGCA) (annealing on the switch adaptor) and a mouse IgH reverse primer mix (mIgG12_r2 (ATTGGGCAGC CCTGATTAGTGGATAGACMGATG), mIgG3_r2 (ATTGGGCAGCCCTGATTAAGGGATAGA CAGATG), mIgA_r2 (ATTGGGCAGCCCTGATTTCAGTGGGTAGATGGTG) binding to the constant region of certain Ig heavy chains). The first PCR was performed with Q5 Hot Start High-Fidelity DNA polymerase (NEB) in a 50µL reaction volume using 7.5 µL of cDNA product with following PCR parameters: 1 cycle of 95°C for 1 min 30 sec; 20 cycles of 95°C for 10 sec, 60°C for 20 sec, 72°C for 40 sec; 1 cycle of 72°C for 4 min; storage at 4°C. PCR 1 products were purified with MinElute PCR purification Kit (Qiagen) and eluted in 25 µL 70°C nuclease-free H_2_O (Qiagen).

Within the second PCR amplification^48^ a M1S primer ((N)4– 6(XXXXX)CAGTGGTATCAACGCAGAG) (annealing on M1SS) and a step-out primer Z ((N)4-6(XXXXX)ATTGGGCAGCCCTGATT), both with sample barcodes, were used. PCR 2 was performed with Q5 Hot Start High-Fidelity DNA polymerase (NEB) in a 50µL reaction volume using 2 µL of PCR 1 product with following PCR parameters: 1 cycle of 95°C for 1 min 30 sec; 14-15 cycles of 95°C for 10 sec, 60°C for 20 sec, 72°C for 40 sec; 1 cycle of 72°C for 4 min; storage at 4°C. PCR 2 products were purified with MinElute PCR purification Kit (Qiagen) and eluted in 25 µL 70°C nuclease-free H_2_O (Qiagen). The products were also gel-purified from 2% agarose gels (extraction with MinElute gel extraction Kit (Qiagen); elution in 15 µL 70°C nuclease-free H_2_O (Qiagen).

Adapter ligation was performed using the TruSeq® DNA PCR-Free Library Prep protocol (Illumina). The products were gel-purified from 2% agarose gels instead of bead purification as mentioned in the protocol (extraction with MinElute gel extraction Kit (Qiagen); elution in 10 µL 70°C nuclease-free H2O (Qiagen)).

The quality of amplified libraries was verified using an Agilent 2100 Bioanalyzer (2100 expert High Sensitivity DNA Assay). According to the fragment size, the libraries were quantified by qPCR using the KAPA Library Quantification Kit for Illumina platforms (KAPA Biosystems).

Based on the result of the qPCR a final library pool with a concentration of 2 pM was used for sequencing with NextSeq500/550 (Illumina) using the NextSeq® 500/550 Mid Output Kit v2 (150 cycles) for 2×150bp paired-end sequencing with 20% PhiX-library spike-in.

Ig repertoire analysis was performed using MIGEC-1.2.4a^49^ in default parameter settings while adding a demultiplexing step for identification of IgG1/2, IgG_3_ and IgA heavy chains. After the MIGEC pipeline’s “checkout” step isotypes were classified according to the presence of mIgG12_r2, mIgG3_r2 and mIgA_r2 primer sequences: AGTGGATAGACMGATG, AAGGGATAGACAGATG and TCAGTGGGTAGATGGTG, allowing for one mismatch against the primer sequence. Data were then processed independently for each isotype. The MIGEC segments file was adjusted to include only C57BL/6-specific V genes for mapping. MIGEC performs a UMI-guided correction to remove PCR as well as sequencing bias and errors. Each resulting consensus sequence was treated as one clone. Clones with identical V, D, and J gene compositions and CDR3 nucleotide sequences were defined as clonotypes. Solely clonotypes consistently found in both technical replicates of a given sample were considered in downstream analyses. Statistics on the overlap of repertoires between different samples were performed based on the presence of clonotypes. To assess likelihood of the observed presence of clonotypes exclusively in one organ or an overlapping presence in both being the result of differences in clone numbers or rare clones at purely random distribution, clones were randomly distributed 1000 times among the samples. Random distributions of sequences to paired samples were drawn while retaining the initial samples’ numbers of clones. Overlap statistics represent the median overlap and clonotypes exclusive to one sample from the 1000 randomizations. The degree of similarity between samples accounting for the abundance of clonotypes is represented by the cosine similarity^50^.

Ig repertoire data discussed in this publication are available at GEO under accession number XXX (Link).

### Statistics and data representation

Absolute numbers of mouse B cell subsets per organ were calculated based on their frequency in a sample. For spleen, peripheral and mesenteric lymph nodes and Peyer’s patches, total organs were prepared and the total numbers of B cell populations calculated based on the numbers in a defined volume determined by flow cytometry (MACSQuant, Miltenyi). BM cell numbers were calculated analogously based on cell numbers in a single femur of a mouse which is estimated to harbor 6.3% of total BM leading to a conversion factor of 7.9 from two femurs to total murine BM^6^.

Further analyses and statistical tests were performed within the R programming environment^51^, with use of the non-base package VennDiagram^52^. Whiskers in Tukey boxplots span 1.5IQR.

### Preparation of histological sections and confocal microscopy

Spleens and femoral bones were fixed in 4% Paraformaldehyde (Electron Microscopy Sciences) for 4h at 4°C, equilibrated in 30% sucrose/PBS, then frozen and stored at -80°C. 6µm cryosections were stained for 1h in 0.1% Tween-20 (Sigma-Aldrich)/10% FCS/PBS after blocking with 10% FCS/PBS for 1h. The following primary and secondary reagents were used: anti-mouse IgG2b-AlexaFluor546 (RMG2b-1, BioLegend); anti-GFP-AlexaFluor488 (rabbit polyclonal, Life Technologies; rat monoclonal FM264G, BioLegend), anti-fibronectin (rabbit polyclonal, Sigma Aldrich), biotinylated anti-mouse Ki67 (Sol-15, eBioscience), biotinylated anti-mouse VCAM-1 (429, eBioScience), anti-human/mouse cadherin 17 (rabbit polyclonal, R&D), anti-mouse laminin (rabbit polyclonal, Sigma Aldrich), anti-mouse IgD-AlexaFluor647 (11.26c, DRFZ), anti-mouse Thy1-Alexa Fluor 594 (T24, DRFZ), anti-mouse B220-AlexaFluor594/647 (RA3.6B4, DRFZ), anti-mouse CD11c-AlexaFluor647 (N418, DRFZ), donkey anti-rabbit polyclonal IgG-AlexaFluor488/647 (Thermo Fisher), strepatavidin-AlexaFluor594/647 (Thermo Fisher). For nuclear staining, sections were stained with 1 μg per ml DAPI in PBS. Sections were mounted in Fluorescent Mounting Medium (DAKO). For confocal microscopy, a Zeiss LSM710 with a 20×/0.8 numerical aperture objective lens was used. Images were generated by tile-scans and maximum intensity projection of 3-5 Z-stacks each with 1μm thickness. Image acquisition was performed using Zen 2010 Version 6.0 and images were analyzed by Zen 2012 Light Edition software (Carl Zeiss MicroImaging). Co-localization of Bsm with cells expressing other markers was performed using sections from three biological replicates. More than 40 cells were inspected per analysis.

### Manual image analysis

Nearest neighbors of Bsm were enumerated based on either direct cell-cell contact or a position within a 10µm radius of cell boundaries of Bsm using high resolution images acquired by confocal microscopy. Cells of interest were identified by immunofluorescent staining for the marker in question. Cells were counted as neighboring cells if pixels from both cells were in either direct contact or within the 10µm radius of cell boundaries.

### Modelling random co-localization

To determine the probability of Bsm cells randomly co-localizing to their observed nearest neighbors, we employed modelling of random cell positioning, in a modification of the previously published approach^7^. In brief, images of Bsm were positioned on histological images of BM at random, repeatedly. Frequencies of co-localizing Bsm and stromal cells were then determined and compared to the frequencies of the original histological images.

## Acknowledgements

We thank Tuula Geske, Heidi Hecker-Kia and Heidi Schliemann for expert technical help, Toralf Kaiser and Jenny Kirsch for support in cell sorting, and Manuela Ohde, Vivien Theißig and Patrick Thiemann for animal care. We thank Sven Künzel and Christine Pfeifle, MPI for Evolutionary Biology in Plön, for providing help and expertise on wild mice. We would like to thank Dr. Dmitriy M. Chudakov for kindly providing the protocol for the generation of libraries for BCR receptor repertoire sequencing. This work was supported by Deutsche Forschungsgemeinschaft through DFG priority program 1468 IMMUNOBONE (to AR and HDC) and the TR130 (to AR and HDC), and by the European Research Council through the Advanced Grant IMMEMO (ERC-2010-AdG.20100317 Grant 268978 to AR). AEH was funded by the DFG (TRR130, HA5354/6-1 and HA5354/8-1). The work of RR and DS was funded in part by the Berlin-Brandenburg School for Regenerative Therapies. RA is a member of Berlin-Brandenburg School for Regenerative Therapies. JZ was a member of GRK1121 (ZIBI) and MW, KW, and SD were members of International Max Planck Research School for Infectious Diseases and Immunology Berlin (IMPRS-IDI). MFM is supported by e:Bio Innovationswettbewerb Systembiologie, a program of the Federal Ministry of Education. Work was supported by the state of Berlin and the “European Regional Development Fund” to G.A.H., F.H., P.M, M.M. and M.F.M. (ERDF 2014–2020, EFRE 1.8/11, Deutsches Rheuma-Forschungszentrum).). HDC is funded by the Dr. Rolf M. Schwiete Foundation. The work of VG, UM and STR work was funded in part by the Swiss National Science Foundation (Project #: 31003A_143869, 31003A_170110 to STR), SystemsX.ch – AntibodyX RTD project (to STR), Swiss Vaccine Research Institute (to STR). PM is supported by EUTRAIN, a FP7 Marie Curie Initial Training Network for Early Stage Researchers funded by the European Union. The DRFZ is a Leibniz Institute. The authors have no conflicting financial interests.

---

**Figure S1:**
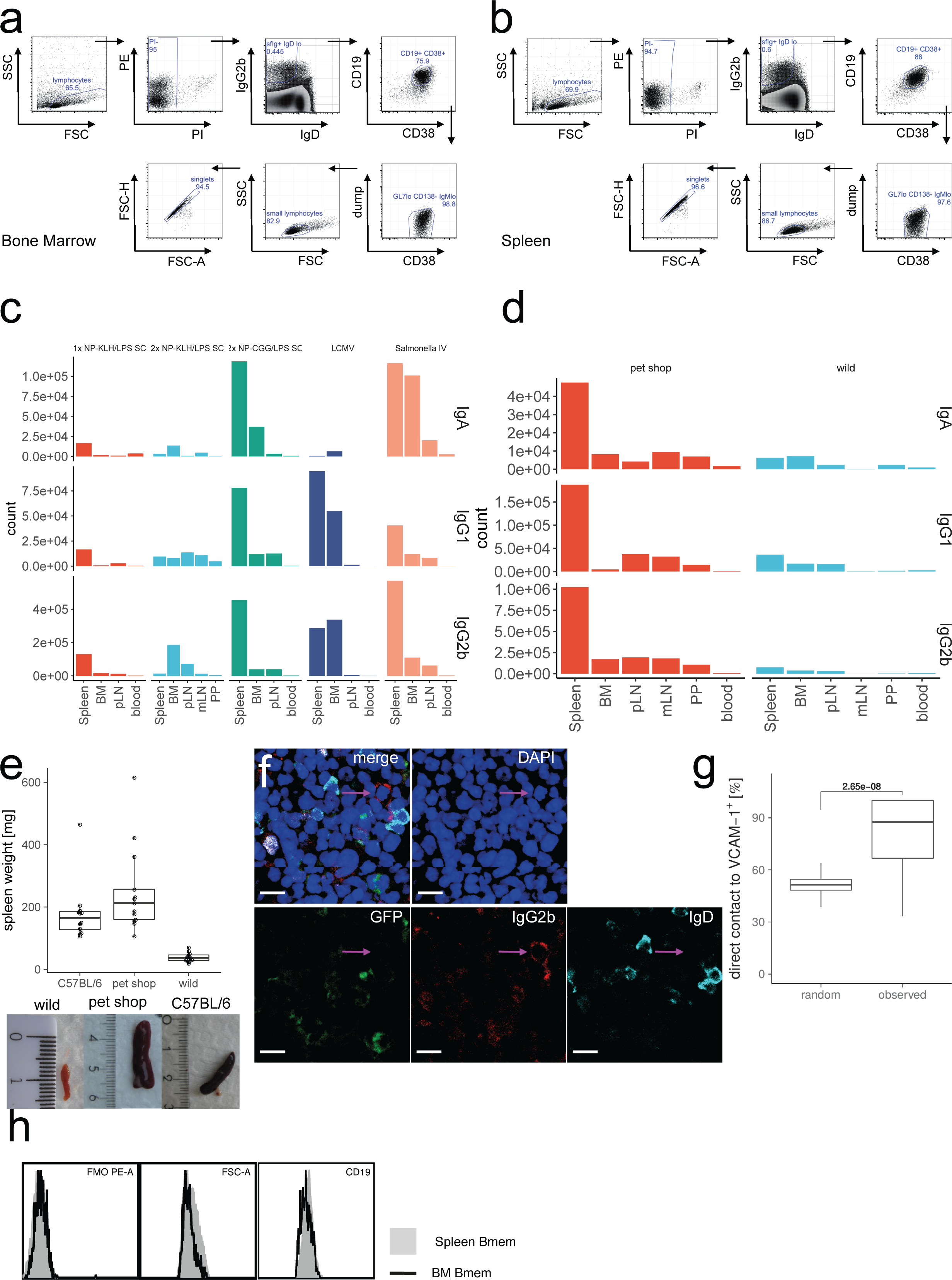
Gating for isotype-switched memory B cells of bone marrow and spleen. BM (a) or spleen (b) CD19+ cells were isolated by MACS technology and switched-memory B cells were identified by expression of surface IgG2b, IgG1, or IgA and CD19, CD38 and lack of IgD, IgM, CD138 and GL7 marker. Staining shown for IgG2b exemplarily. c) Cell numbers of Ig-switched memory B cells per organ in C57BL/6 laboratory mice. Absolute cell numbers per organ calculated from flow cytometric counts (gated for IgG1+, IgG2b+, or IgA+ CD19+CD38+CD138-GL7-CD11c-IgM-IgD-PI-small lymphocytes); n=42, data pooled from 8 experiments with five different immunizations protocols performed in C57BL/6 mice aged 4-20 months and held under SPF conditions, data presented as median cell count per organ by immunization. BM: bone marrow, pLN: peripheral lymph nodes, mLN: mesenteric lymph nodes, PP: Peyer’s patches. d) Cell numbers of Bsm per organ in wild and pet shop mice. Median cell numbers per organ calculated from flow cytometric counts (gated for IgG1+, IgG2b+, or IgA+ CD19+CD38+CD138-GL7-CD11c-IgM-IgD-PI-small lymphocytes); n=13 (wild) and 13 (pet). e) Spleen weight from spleens of C57BL/6 mice held under SPF conditions, pet shop and wild mice, n=10 (C57BL/6), 13 (pet), and 11 (wild). f) Identification of bone marrow IgG2b+ Bsm. Naive B cells and plasma cells were excluded by IgD staining and Blimp1-GFP signal, respectively. Cell nucleus was identified with DAPI (blue). Scale bar: 10µm. g) Simulation of co-localization between Bsm and VCAM-1+ stromal cells. Non-random co-localization of bone marrow Bsm was determined using images acquired from 7 bone marrow slides. Graphs represent direct co-localization of more than 12000 simulated cells (random) versus co-localization observed per slide for 28 slides from 4 mice with two or more analyzed Bsm per slide (mean=5.66 cells per slide), p value (Welch’s test) indicated on graph. h) Control stainings for spleen and bone marrow (BM) memory B cells in flow cytometry, identified as IgG2b+CD19+CD38+CD138-GL7-CD11c-IgM-IgD-PI-small lymphocytes.

**Figure S2:**
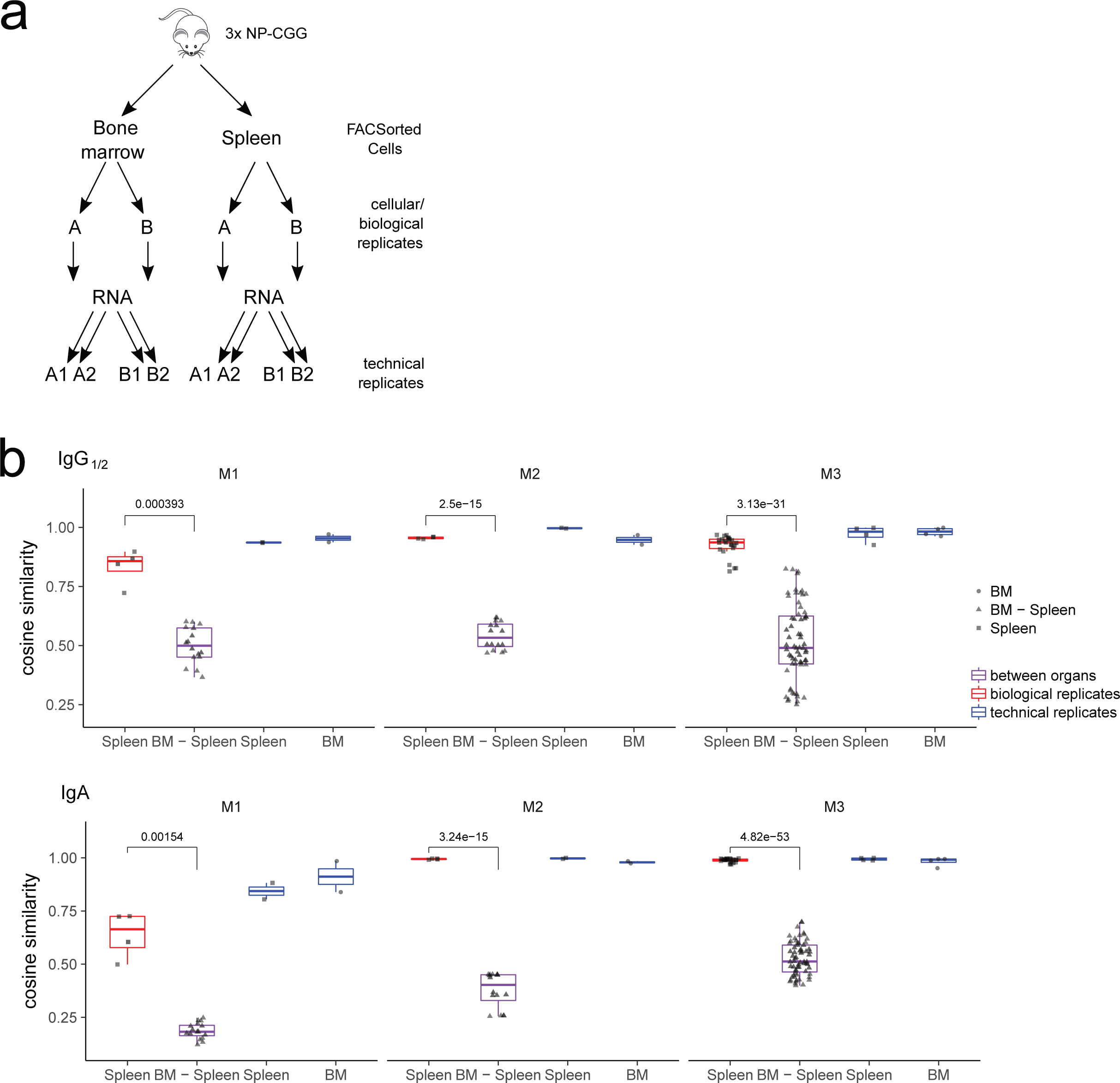
a) Experimental setup for the comparison of repertoire of isotype-switched memory B cells of BM and spleen: after isolation, cells of the same organ were divided into equal proportions and processed as biological replicates. After RNA isolation, samples were split and processed as technical replicates. b) Cosine similarity between samples of IgG1/2+ (upper panel) and IgA+ (lower panel) heavy chain CDR3 repertoires accounting for clonotype frequencies. Graphs represent cosine similarity comparison within technical replicates of spleen and BM (blue), within cellular replicates from spleens (red), and between spleen and bone marrow (BM-Spleen, purple) of three individual mice. p values (Welch’s test for difference of means of cosine similarity within shared IgH repertoire (spleen cellular replicates) and between spleen and bone marrow replicates are indicated.

**Figure S3:**
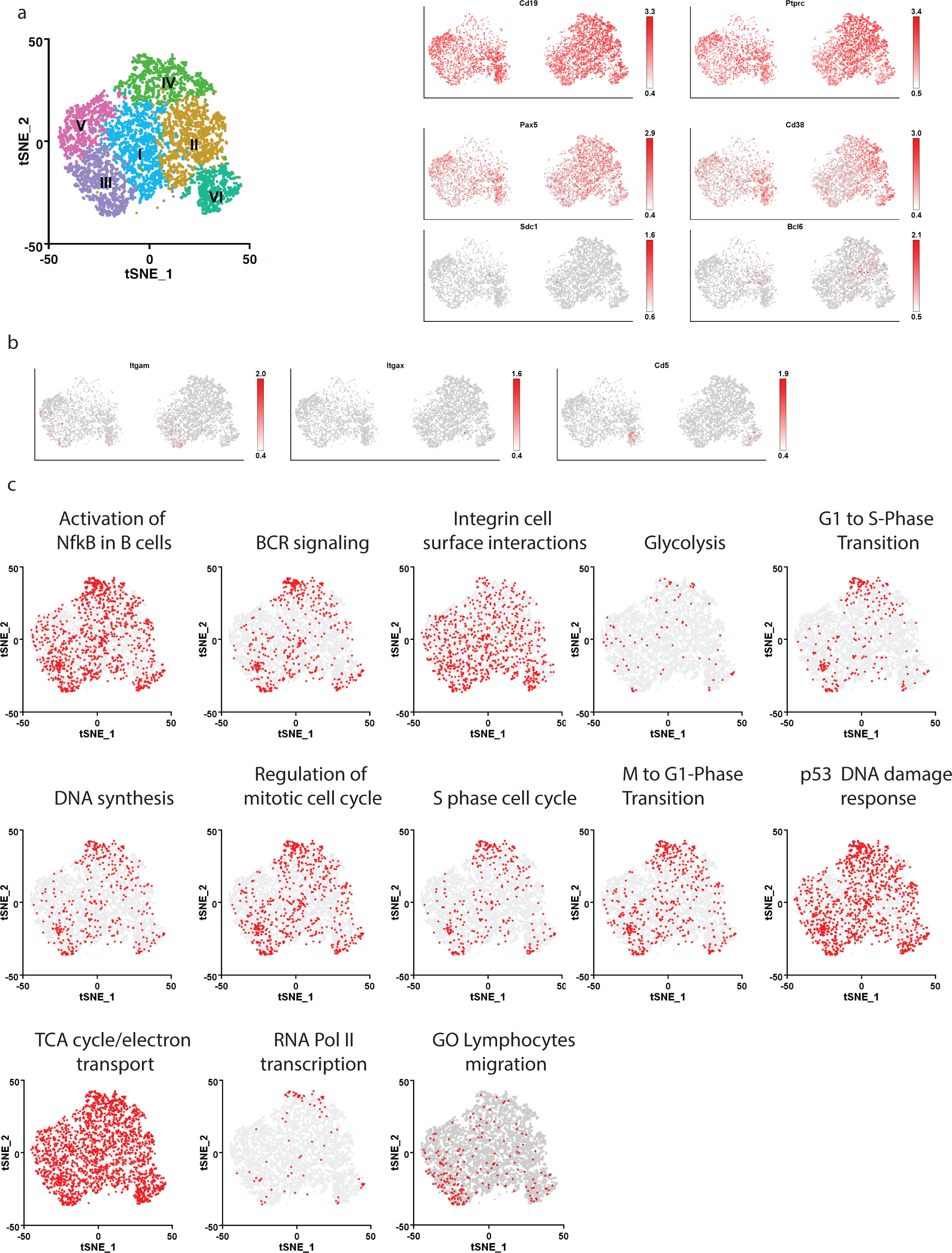
a) Six transcriptionally defined Clusters were identified by shared nearest neighbor (SNN) modularity optimization based clustering algorithm mapped to tSNE representation of spleen and bone marrow (BM) cells. tSNE coordinates and clustering was computed for 4754 from spleen and 2947 from BM cells, representation combined for spleen and BM (left). Distribution of expression for *Cd19*, *Ptprc* (CD45), *Pax5*, *Cd38*, *Sdc1* (CD138), and *Bcl6* genes mapped on tSNE representation. b) Distribution of expression for *Itgax* (CD11c), *Itgam* (CD11b), *Cd5* genes mapped on tSNE representation. c) Single cell gene set enrichment analysis of expression of the Reactome and Gene ontology enrichment (GO) mapped on tSNE representation. Cells for single cell sequencing were FACSorted as IgG-expressing CD19+CD38+CD138-GL7-small lymphocytes.

## Abbreviations

Bsm: isotype-switched memory B cell
BM: bone marrow
CGG: chicken γ globulin
IFA: incomplete Freund’s adjuvant
i.p.: intraperitoneal
KLH: keyhole limpet hemocyanin
LCMV: lymphocytic choriomeningitis virus
LPS: lipopolysaccharide
MANOVA: Multivariate analysis of variance
NP: 4-hydroxy-3-nitrophenylacetyl
s.c.: subcutaneous
SPF: specific pathogen-free
TT: tetanus toxoid

